# Closed-loop coupling of a muscle to a robotic device for dynamic assessment of muscle function

**DOI:** 10.1101/337303

**Authors:** Kartik Sundar, Lena H. Ting, Stephen P. DeWeerth

**Author notes:** Corresponding Author: Stephen P. DeWeerth, The Wallace H. Coulter Department of Biomedical Engineering, Georgia Tech and Emory University, 313 Ferst Dr., Atlanta GA 30332.

## Abstract

The contributions of individual muscles to the performance of functional tasks are difficult to evaluate using traditional isolated muscle protocols. During movements, skeletal muscles work against a variety of environmental loads that influence their energetics and function. In turn, these changes in muscle length and muscle velocity alter the forces that the muscle can generate. Classic single-muscle experiments clamp at least one muscle state (length, velocity, force) such that it is independent of the other states, interrupting the dynamic interactions between the muscle and its environment. The purpose of this study was to design and build a real-time feedback system to virtually couple an isolated muscle to a robotic device. Using this approach, the muscle length is not prescribed, but results from the dynamic interactions between the muscle and a physical environment. Therefore our device facilitates the study of how physical interactions between a muscle, limb, and environments alter the force and motion produced by the muscle during controlled muscle activation. To demonstrate the utility of our system, we replicated some salient features of frog swimming, we coupled a frog plantaris longus muscle to a one-degree of freedom “limb” that drove a frog foot through water. We demonstrate that under identical muscle stimulation parameters, changes to muscle moment arm, environmental viscosity, and muscle fatigue can significantly alter the resulting muscle force, length, and work.

## I. Introduction

Animal locomotion arises from complex nonlinear interactions between the neuromuscular system and its natural environment. Quantifying the mechanical properties of a muscle as it interacts with the environment is essential to understanding the strategies that underlie movement. Muscle function is difficult to quantify in behaving animals because experimental manipulation and measurements of quantities such as force and length are challenging to achieve. In contrast, detached or isolated muscle preparations facilitate controllability and high-resolution data collection but do not replicate the interactions between the muscle and the natural environment. Further, isolated muscle preparations are typically designed to examine a specific muscular component and do not reveal the synergetic effects of the different components working together. By virtually connecting an isolated muscle to a physical robotic device, we introduce a closed-loop neuromechanical system to study muscle properties during functional dynamic conditions where muscular and environmental forces interact to produce motion.

*In vivo* muscle measurement techniques allow individual muscle function to be studied during the complex interactions between the neuromuscular system and the environment during natural movement conditions. For example, EMG electrodes, sonomicrometry crystals, and buckle tendon transducers can be used to describe changes in muscle length, force, and activation during locomotion and can be used to examine whole-muscle dynamics in an intact animal during locomotion [1-3]. However, *in vivo* methods only allow the examination of one or a few muscles amongst many that contribute to the movement. Therefore it is difficult to ascertain the precise role of a particular muscle to the overall motion of the body. Further, experimental manipulations to test hypotheses about single muscle function are difficult to impossible *in vivo* [4]. Moreover, even when manipulations of a limb or muscle can be introduced, adaptations at both the neural and mechanical level can extend across multiple muscles, limiting the conclusions that can be drawn about the contributions of a single muscle to the production of a movement.

Conversely, isolated *in vitro* muscle experiments are designed to allow the identification of state-dependent muscle properties by explicitly removing the dynamic interactions that vary a muscle’s state during a natural movement [5, 6]. Typically, isolated or detached muscle protocols explicitly specify at least one muscle state (length [7, 8], velocity [9, 10], force [11, 12]) such that it is independent of the other states. For example, single muscle energetics have been measured using the classical work-loop method in which the muscle length is prescribed to move along a sinusoidal path that is independent of muscle force produced through electrical stimulation of the muscle [13-15]. Typically, the muscle is stimulated at different phases of the sinusoidal motion, or for varying durations and the resulting energetics are measured. Such protocols allow muscle properties to be studied under a variety of conditions where particular variables such as muscle length, velocity, or force are controlled well. While these procedure can replicate the particular trajectory and force combinations measured *in vivo*, they cannot provide information about what happens to the movement, and thus the muscle state, if the muscle’s force-producing capabilities should change, such as due to altered stimulation patterns or fatigue, or a change in the load. Thus, in such clamped conditions, because the dynamic interactions between the muscle and its environment are interrupted, the derived muscle properties may differ from those that might be observed under behavioral conditions.

Alternately, closed-loop methods that couple a muscle to a simulated mechanical environment do not require any muscle states to be explicitly specified. Isolated muscle systems using real-time feedback to allow a muscle’s force to move a simulated mechanical load [16-18]. In these systems, the interactions between the muscle and the simulated environment are defined by physical laws of motion such that none of the muscle states have to be predetermined. Rather, movement arises from realistic dynamic interactions that occur as a result of muscle stimulation, force production and the effect of the force on the load. Such protocols have revealed interesting dynamic properties of muscle [16, 17]. However, they are somewhat limited by the ability to adequately simulate the complexity of the natural environment which is often too difficult to model computationally, especially under real-time constraints.

In cases of complex mechanical dynamics, a physical or robotic model of a system can more realistically simulate the salient dynamics of a system than a computational model. Robots or other mechanical models are often used to create and study the complex interactions that occur during locomotion such as fluid dynamics or ground contact [19, 20]. For example, during frog swimming, the load on the muscular system is a function of the viscous resistance of the water on the foot and is complicated by the biomechanics of the frog leg. During the power stroke, the webbed toes open to increase resistance and create forward thrust. During recoil, the webbing closes allowing the leg to move through the water without substantially propelling the frog backwards. A physical model of these interactions would provide a realism that a computational model could not.

The purpose of this study was to develop a *closed-loop neuromechanical system* that applies real-time control to couple an isolated muscle to a physical environment using a robotic device‥ The system allowed us to examine the resulting movement produced when the muscle was stimulated while coupled to different environments, as well as under conditions of muscle fatigue. To illustrate the benefits of the closed-loop neuromechanical system we implemented a simple example of frog swimming. Specifically, we simulated salient features of frog swimming, by virtually coupling a muscle to a single-degree-of-freedom robotic limb immersed in a container of water with a real frog foot attached on the end. We conducted three illustrative experiments to demonstrate how our system enables the study of single-muscle function during a variety of tasks that would be difficult to reproduce using *in vivo* or isolated muscle techniques, and demonstrate how the muscle force, work, and the resulting movement vary as the interactions with the environment change. Under identical muscle stimulation conditions we show that muscle moment arm, environment viscosity, and muscle fatigue can dramatically alter the work produced in a single propulsive stroke. Our approach may therefore facilitate better predictions about neuromuscular strategies and muscle function during complex movements.

## II. SYSTEM ARCHITECTURE AND EXPERIMENTAL DESIGN

The closed-loop neuromechanical system used real-time feedback to couple an isolated muscle and a robotic device (Fig. 1). The architecture, implemented on a real-time processor managed a variety of actuators and sensors in a closed-loop feedback paradigm:

**Figure 1:**
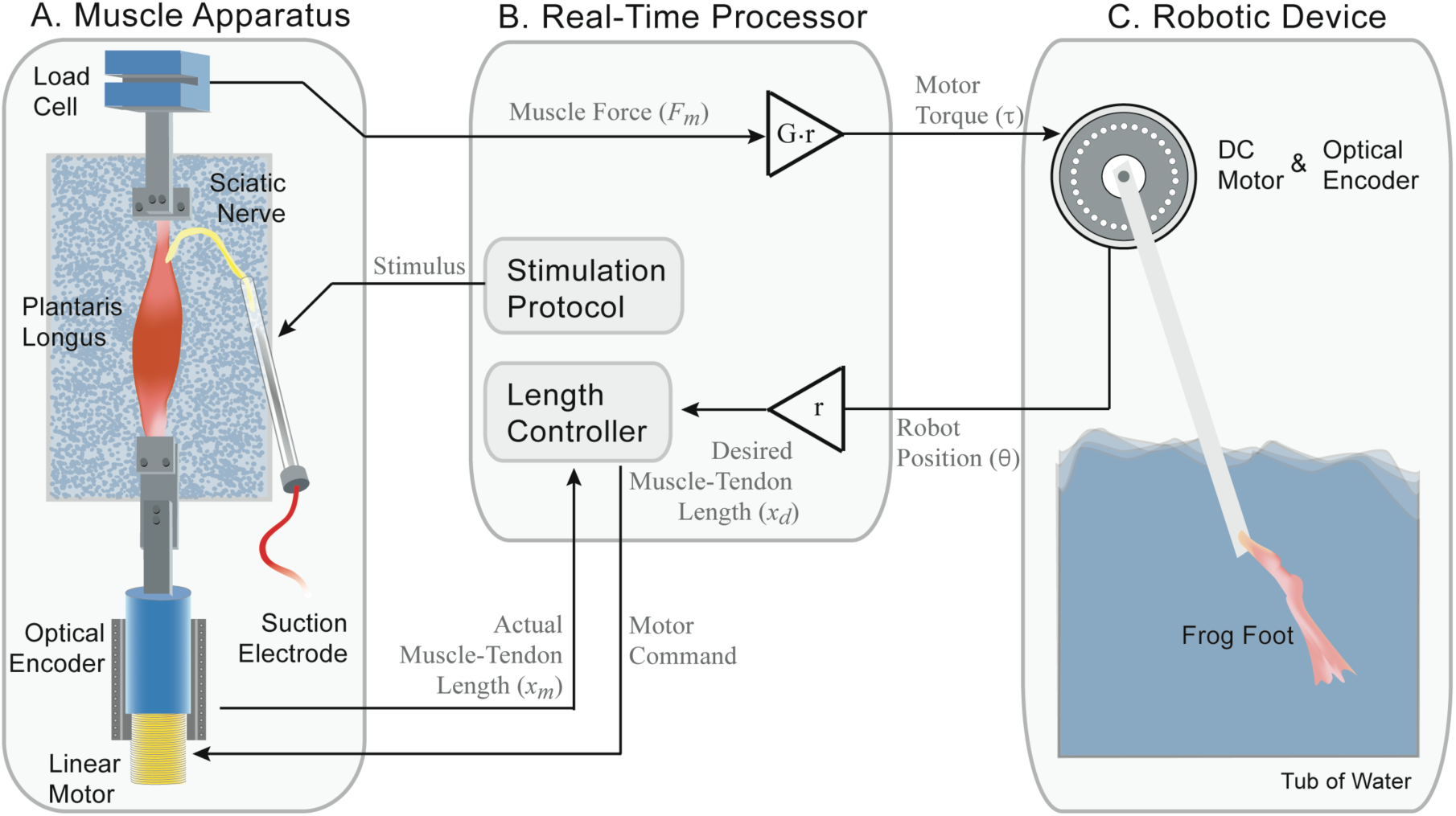
Architecture of the closed-loop neuromechanical system. An isolated muscle (A) is stimulated and a load cell measures the force. The force is transformed by a virtual mechanical model (in this example, a moment arm transformation) running on the real-time processor (B). The resultant torque is generated by a motor in the robotic device (C). The position of the robotic limb (θ) is transformed into a muscle-tendon length (*x*_*m*_). A closed-loop length controller ensures that difference between the actual muscle length and desired muscle length (*x*_*d*_) is minimal.

1. Electrical stimulation activated the muscle, producing a force.
2. The force produced by the muscle was measured and used to specify the torque applied to the robotic device via a torque motor. The robotic device moved according to the forces acted on by both the motor and the external environment.
3. The resulting position of the robotic device was measured and specified the desired length of the muscle-tendon unit, thus closing the loop. A muscle length controller minimized the difference between the actual and desired muscle length.

### A. Design Criteria

The closed-loop neuromechanical system was implemented with a *frog (Rana pipiens*) muscle. The actuators, and sensors used by the closed-loop system exceeded the following specifications:

### Muscle Apparatus

1. The muscle length controller was required to have a steady-state stiffness greater than 80 kN/m, corresponding to a strain of no greater than 1% at maximum isometric muscle force for frog muscles [21].
2. Most mechanical systems the muscle would interact with would have relatively low natural frequencies (0-10 Hz). As such, our system was required to have a bandwidth of a least 25Hz. A relatively flat amplitude response was required such that the controller did not add or remove energy from the system. Changes in amplitude less than 2 dB were considered appropriate. In addition, the phase delays no greater than 10° were also required.
3. A resolution of 10 μm (1% of 1 mm) was required of the muscle position sensor because *in vivo*, the length of muscles in the frog hindlimb can change on the order or millimeters during swimming or jumping [22, 23].
4. Typically, forces in the frog hindlimb range from 1 to 15 N [21]. The muscle force sensor was required to discern changes in muscle force of as small as 1 mN or less.

### Robotic Device

5) The inertia and friction of the torque motor were considered part robotic device. Therefore, we did not require the closed-loop system to compensate for the dynamics of the torque motor. This is a limitation of our system that would need to be addressed for accurate hypothesis testing.

6) Muscle moment arms in the frog hindlimb are on the order of millimeters. Assuming moment arms no greater than 1 cm, the robot position sensor was required to have a minimum resolution of 1000 ticks per radian. This mapped to a resolution of 10 μm for the muscle length.

### B. Muscle Apparatus

We used the isolated frog plantaris longus (PL) to demonstrate the abilities of the neuromechanical system, as the mechanical and energetic properties of frog hindlimb muscles in traditional behavioral and single-muscle preparations are well-known and serve as a good point of comparison [11, 24, 25]. The typical burst activation pattern duration of the PL during motor task can be as long as 150 ms and the muscle can shorten 10% of resting length [22, 26].

All surgeries were performed according to procedures approved by the Institutional Animal Care and Use Committee at the Georgia Institute of Technology (Protocol #A04010). Prior to surgery, one frog (*Rana pipiens*) was anestheized with tricaine methanesulfonate (MS-222, 1 g L^−1^) and then double pithed. The PL, still innervated by its nerve, was removed along with a portion of the sciatic nerve. A bone chip was left at the proximal end and a large piece of tendon (approximately 80-90% of *in vivo* length) was left at the distal end. Small plastic clamps were used to attach the distal tendon to a load cell (Strain Measurement Devices S251) and the proximal bone chip end to a linear actuator (H2W Technologies). The entire muscle was submerged in a bath (22 °C) of oxygenated Ringer solution (NaCl, KCl, CaCl_2_, NaHC0_3_).

A suction electrode was used to stimulate the sciatic nerve and to elicit a force from the muscle. Muscle force (*F*_*m*_) was measured using the load cell. The muscle-tendon length (*x*_*m*_) was controlled using the linear actuator, and the actual muscle-tendon length was measured using a 1 µm resolution optical encoder (Renishaw RGH41X30D05A), exceeding the 10 µm requirement.

### C. Robotic Device

We used a single-degree-of-freedom robotic device consisting of a 0.4 cm diameter, 15 cm length aluminum rod with a frog foot attached on the end (Fig. 2, Table I). The frog foot was cut at the elongated tarsals and rigidly clamped to the device at the tarsometatarsal joint, leaving the webbed toes intact. The robotic device was then driven by a DC torque motor (Faulhaber 2342-024CR) and moved through a container of tap water. The limb was designed so that the morphology of the frog foot played the largest role in creating viscous resistance during movement. While the natural frequency of the aluminum rod was approximately 2 Hz, the entire device was over-damped when placed in water.

**Figure 2:**
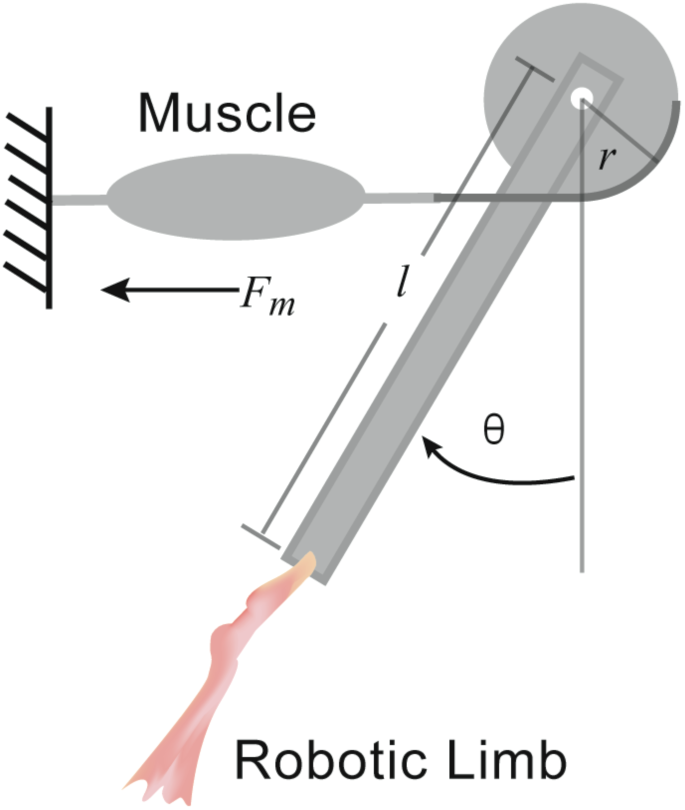
Functional schematic of the closed-loop neuromechanical system. When the system is assembled it functions as a single joint actuated by a one muscle with a constant moment arm. In this configuration muscle force (*F*_*m*_) causes an increase in joint angle (θ). Gravitational and other environmental forces can act to decrease the joint angle. The force produced by the muscle (*F*_*m*_) is amplified by a gain (G) that is not shown in this schematic.

**Table I.** M<u>ECHANICAL</u> QUANTITIES OF THE NEUROMECHANICALSYSTEM

| Symbol | Quantity | Value |
| --- | --- | --- |
| $\theta$ | angle, robotic limb | Output |
| $L$ | length, robotic limb | 15 cm |
| $M$ | mass, robotic limb | 10 g |
| $r$ | length, moment arm | Varied |
| $F_m$ | force, muscle | Output |
| $F_i$ | force, initial muscle-tendon | $\sim 0$ N |
| $G$ | gain, force | 3 |
| $x_m$ | length, actual muscle-tendon | Output |
| $x_d$ | length, desired muscle-tendon | Output |
| $x_i$ | length, initial muscle-tendon | Lo* (20 mm) |

\* Lo is defined as the length where the muscle-tendon unit can produce the maximum isometric force

Torques applied via the DC motor caused the device to rotate, and the position (θ) was measured using an optical encoder (US Digital E3 2500 CPR) which had a resolution of more than 1500 ticks per radian (1000 required). To accelerate the device, the muscle was required to produce enough force to overcome the viscous resistance of the frog foot moving through the water, the inertial forces of the robotic device, gravity, and other nonlinear forces such as friction.

### D. Real-Time Processing

A real-time processor (dSPACE Inc. DS1104) converted the muscle force (*F*_*m*_) to a torque (τ) that was applied to the robotic device using the DC motor. The torque applied to the robotic device was determined by the following equation:

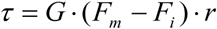

where *r* is the virtual moment arm, *G* is a gain term used to amplify the muscle force. Forces produced by the living muscle were referenced to an initial background force (*F*_*i*_). This allowed the muscle to apply positive and negative changes in force requiring only one muscle to actuate the robotic device in either direction.

Sampled at 10 kHz, the position of the robotic device (θ) was used to determine the muscle-tendon length (*x*_*m*_). The device was connected to the frog muscle via a constant virtual moment arm (*r*). The desired muscle length was computed by the following equation:

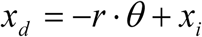

where *x*_*i*_ is the initial muscle-tendon length. A muscle length controller, running on the processor, minimized the difference between the desired length (*x*_*d*_) and the actual length (*x*_*m*_).

### E. Experimental Design

At the start of each experiment trial, the robotic limb was aligned vertically, and the initial muscle-tendon length (*x*_*i*_) was set to the optimal muscle length L_o_ (the length where the muscle is able to produce maximum active isometric force), which was determined experimentally from twitch contractions at various lengths. The optimal length (L_o_) for the PL muscles tested were approximately 20 mm. The PL was maximally activated for 100 ms (approximately equal to the period of PL activity measured in *Rana pipiens* swimming [26]), and the resultant kinematics were measured. The activation was achieved using a stimulus frequency of 200 Hz and a pulse-width of 100 μs. The stimulus current was adjusted until maximum activation was achieved. Between each trial the muscle was allowed to rest for two minutes. Isometric contractions were periodically used to check the viability of the muscle. Muscle fatigue was quantified by the percentage drop in isometric force measured at the optimal length. During the experiments, the muscle was visually inspected, and muscle force recordings were checked to ensure that the muscle did not slip. After data collection was completed, the PL was removed from the bath, all non-muscular tissue was cut away, and the resultant muscle tissue was weighed.

### F. System Validation

The closed-loop architecture ensured that the virtual connection between the muscle and robotic limb closely resembled a real physical connection. Specifications of the actuators and sensors used by the neuromechanical system exceeded the requirements previously described and are listed in Table II.

**Table II.**
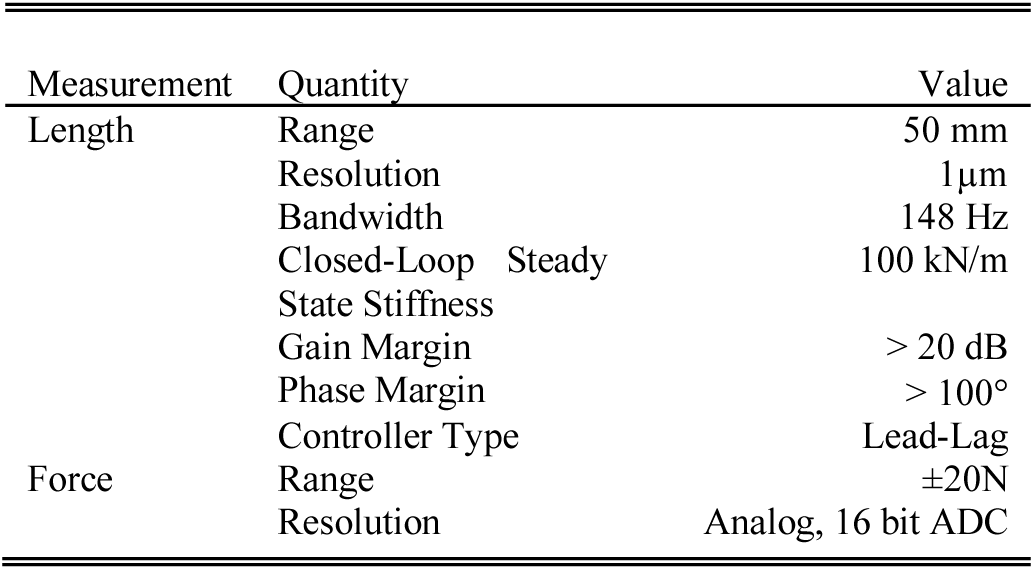
MUSCLE CONTROL SPECIFICATIONS

The muscle length controller was implemented using a second order lead-lag cascade. First, a computational model of the linear actuator was developed and an initial lead-lag controller that met the design criteria was constructed. By trial and error, the initial controller design was tested and modified with the linear actuator in the loop. In addition, the controller was tested with muscles and springs of various compliances to ensure that controller remained stable. The muscle length controller had a steady-state stiffness of approximately 100 kN/m, surpassing the requirement of 80 kN/m. The frequency response of the controller, without a muscle attached, was experimentally determined by sweeping frequencies with a small amplitude of 0.02 mm (Fig. 3). The controller was band-limited at approximately 30 Hz (25 Hz required), which was much greater than the natural frequency of our robotic limb (0-2 Hz). Larger amplitudes saturated the current limit for the muscle length actuator and decreased the bandwidth. During all experiments, the actuator current was monitored to confirm that it was not saturated and that muscle control was not compromised. The muscle length magnitude response was relatively flat (within 2 dB) up to 120 Hz (Fig. 3A). The phase response (Fig. 3B), however, showed that the controller exhibited time delays at frequencies beyond 30Hz.

**Figure 3:**
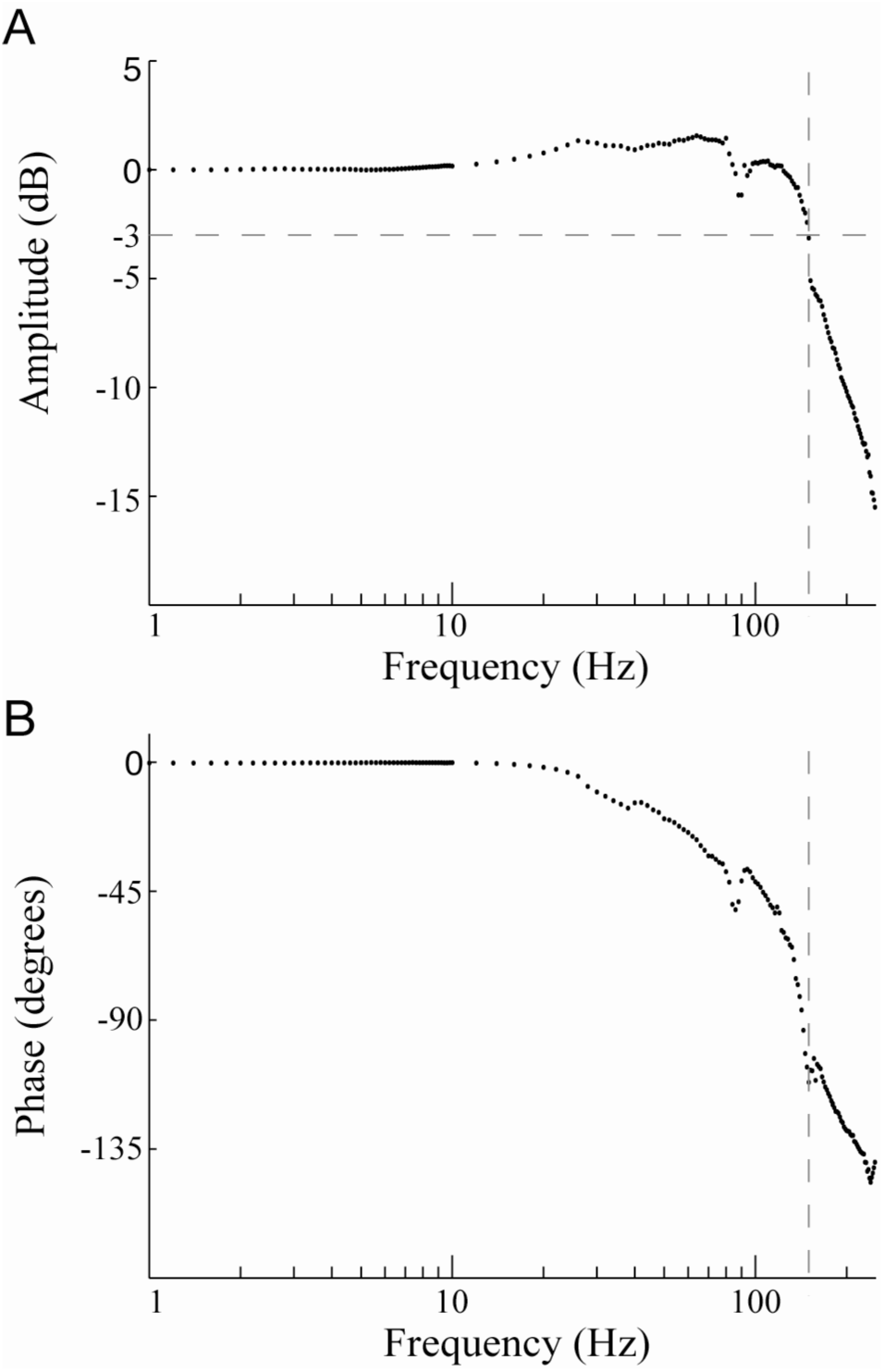
Closed-loop frequency response of the muscle length controller. The magnitude (A) and phase (B) response of the transfer function was experimentally obtained by sweeping the frequency of the desired muscle length (*x*_*d*_) and measuring the actual muscle length (*x*_*m*_). The reference desired muscle length signals (*x*_*d*_) had an amplitude of 0.02 mm. The-3 dB bandwidth was measured to be 148 Hz and within the majority of the bandwidth (0-120 Hz) changes in the magnitude response were less than ±2 dB.

To facilitate the closed-loop validation, we replaced the muscle with a spring and checked to ensure that the stiffness of the spring (4 N/mm) would be effectively emulated to the robotic device. The robotic device was moved and the resultant length and force of the spring was measured – the stiffness measured through the closed-loop system (3.999 N/mm) was equivalent to the actual stiffness of spring. To further validate the closed-loop system, we then replaced the robotic device with a computational mass (similar to [16-18]). After applying a step response to the system, we compared the physical oscillations of the real spring (thick gray line) to that of the completely theoretical spring-mass system (thick grey line, Fig. 4B). The dynamics of the real spring closely matched that of the theoretical spring-mass system. Over the course of the 1 second recording, the amplitude of the oscillation drifted by less than 0.5% due to the frequency response of the muscle length-controller. These errors would be even further reduced if damping were added to the system, as in the case of living muscle or inserting the robotic device in water. We also varied the computational mass and the size of the step response to ensure that the frequency and magnitude of the oscillations changed accordingly. Assuming high accuracy of the position sensors on the robotic device and rapid ability to apply torques, there are no significant differences between the virtual coupling created by our system and a physical coupling within the prescribed parameters (see Design Criteria).

**Figure 4:**
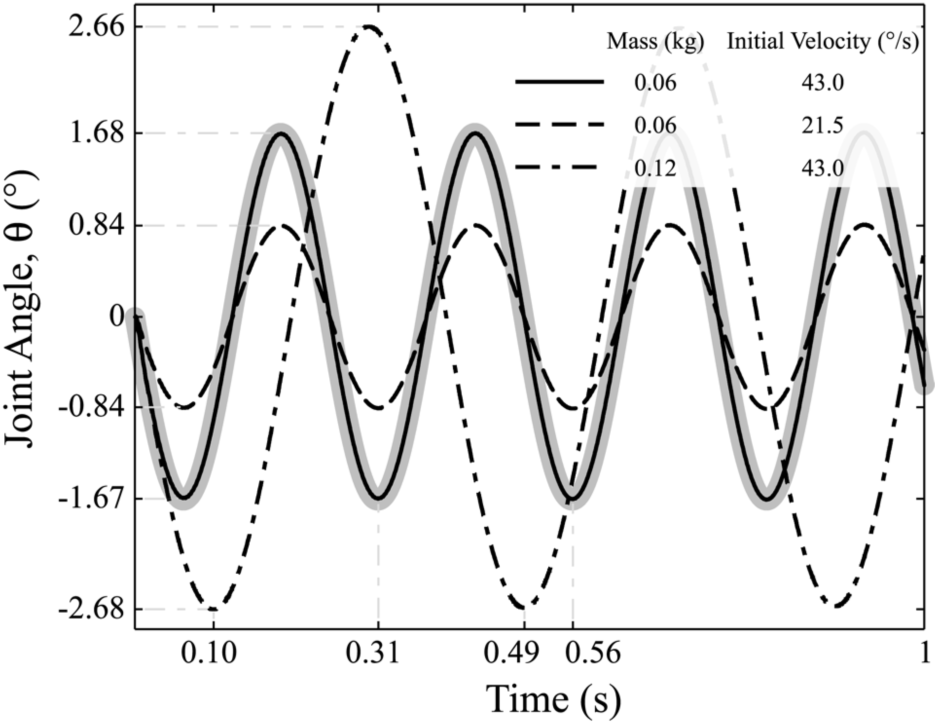
Closed-loop performance of the system with a physical spring in place of the muscle. The dynamics of the physical spring when connected to a computationally-simulated loads. The motion of the physical spring (thick gray line) was compared to simulated spring-mass system (thin gray line) for validation. As expected, the actual trajectory closely matched the theoretical trajectory. The motion of the physical spring when attached to other simulated masses are also shown (dashed lines).

In addition, we compared the torques applied to the robotic device (τ) and the actual muscle length (*x*_*m*_) to their respective desired values during a typical experiment (Fig. 5). The torques applied to the robotic device (converted to Newtons for comparison by dividing by the gain (*G*) and moment arm (*r*)) accurately matched the forces produced by muscle. The maximum error was 3.3 × 10^−5^N and therefore, the performance of the entire system is constrained by the dynamics of the muscle length controller. The actual muscle length also accurately matched the desired muscle length (*x*_*d*_) (which is equivalent to the position of the robotic device (θ)). Because forces produced by the muscle acted to displace the linear actuator, the actual muscle length led the desired muscle length when the muscle was producing a force. Due primarily to load applied to actuator by the muscle, and not the dynamics of the lead-lag controller, the maximum error during a typical experiment was 0.12 mm or 0.6% of optimum length. In addition, 99% of the power of the muscle length movement during a typical experiment was within 0-2 Hz and well within the capabilities of the muscle-length controller.

**Figure 5:**
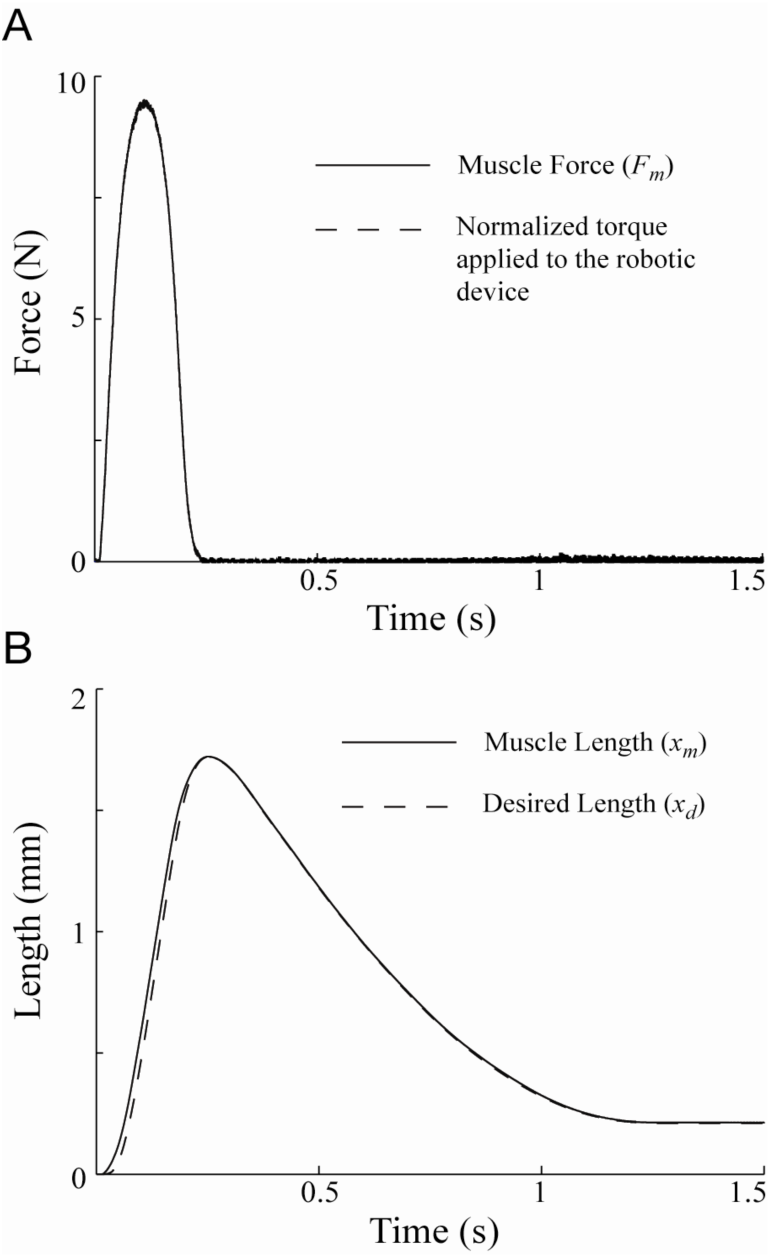
Closed-loop performance of the system during a typical experiment. (A) The force produced by the muscle (*F*_*m*_) and the torque (τ, converted to Newtons for comparison by dividing by the gain (*G*) and moment arm (*r*)) applied to the robotic device were measured and compared. As desired, the two data sets are indistinguishable validating our ability to accurately apply the forces produced by an isolated muscle to a robotic device. (B) The actual muscle length (*x*_*m*_) closely tracked the desired muscle length (*x*_*d*_). Because of the forces imposed on the linear actuator by the muscle, we found that the actual muscle length (*x*_*m*_) led the desired muscle length (*x*_*d*_). Overall, our closed-loop system performed within the limits of the desired criteria.

## III. EXPERIMENTAL VALIDATION

In order to demonstrate the utility and benefits of the closed-loop neuromechanical system, we conducted three example experiments that varied (1) muscle moment arm, (2) environment viscosity, and (3) muscle fatigue. We show how these variations alter the interactions between the muscle and the environment to affect muscle kinematics and energetics under identical muscle stimulation conditions (Fig. 6, Table III). We compare the muscle force, muscle length, as well as the net work produced under the three conditions. These examples illustrate how our approach combines the benefits of current *in vitro* and *in vivo* methods.

**Figure 6:**
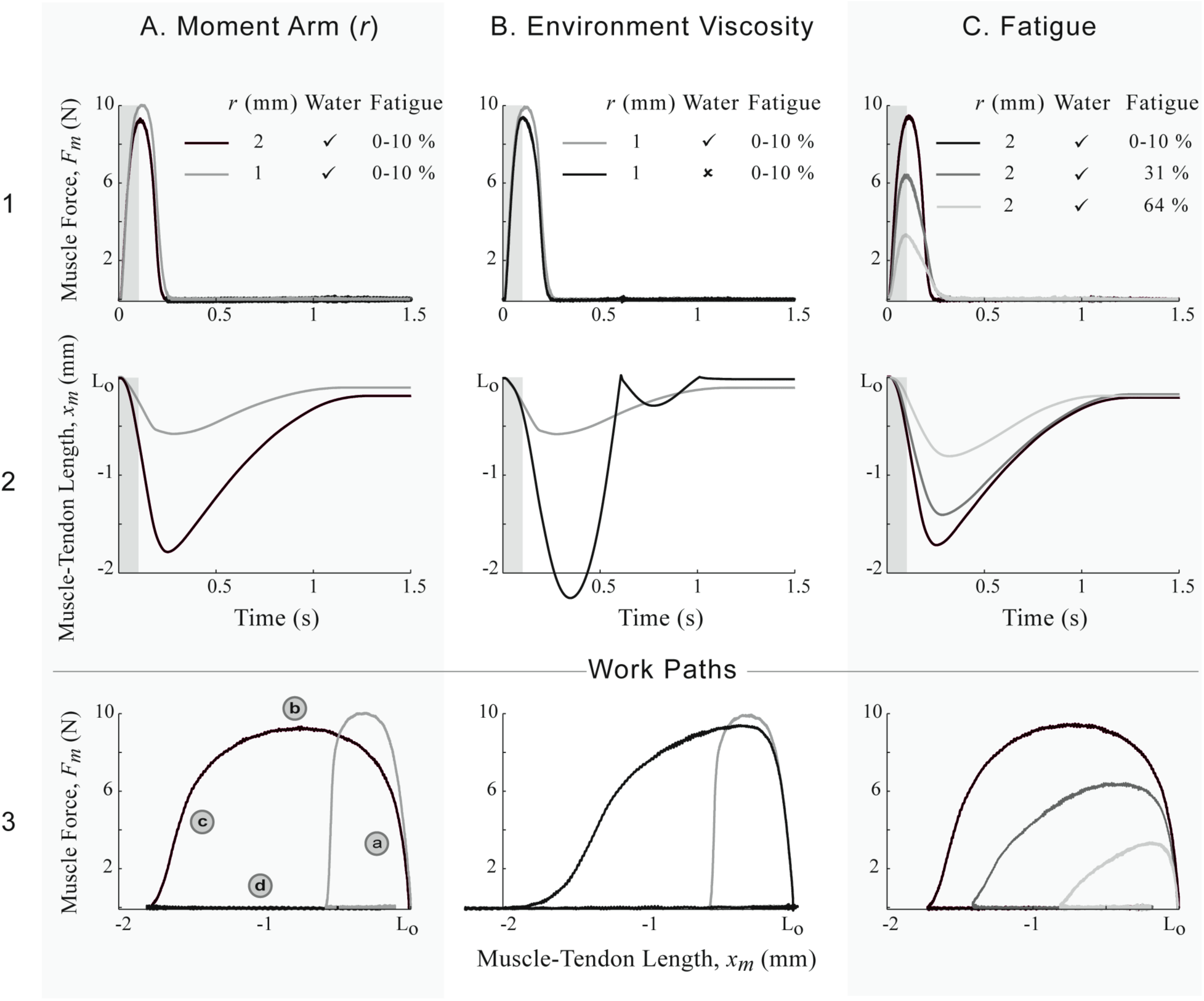
Muscle length and force trajectories. Each column shows the muscle-tendon length, force, and work-loop trajectories for a variation in one system parameter: (A) muscle moment arm, (B) environmental viscosity, and (C) muscle fatigue. Rows 1 and 2 show the muscle force and length time responses respectively. The duration of muscle stimulation is indicated by the shaded rectangle. The third row plots muscle force versus length to demonstrate the work loop for each experiment. The progression around the work loop is shown in the lower left panel with the following four stages: (a) Upon muscle activation, the muscle force rises without substantial shortening. (b) The power stroke is produced when the muscle shortens while producing a large constant force. (c) After the stimulation is stopped, muscle force declines while inertia causes the muscle to continue to shorten. (d) The muscle passively lengthens due to gravity acting on the robotic device.

**Table III.**
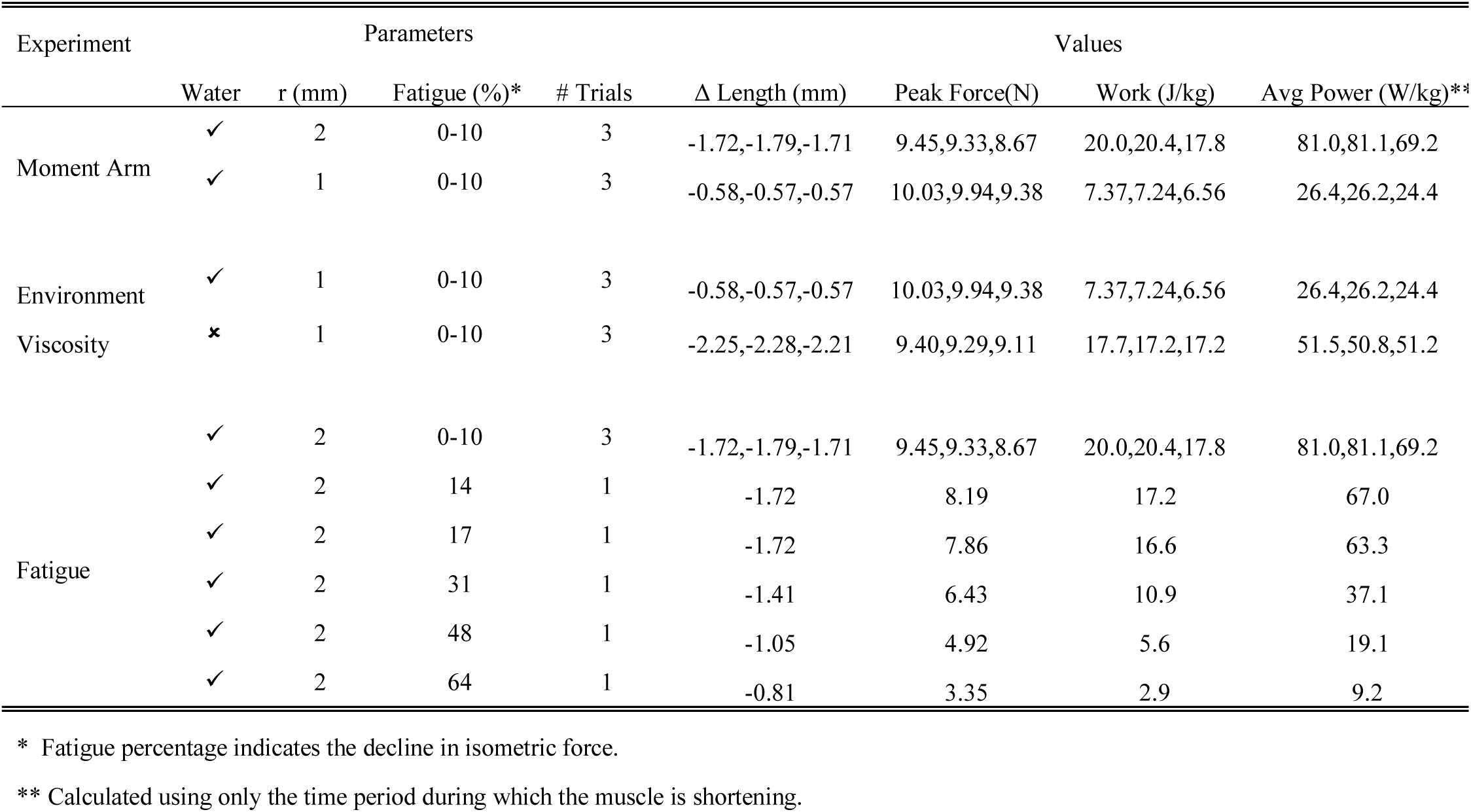
MUSCLE ENERGY AND KINEMATIC MEASUREMENTS

### A. Closed-loop “Swimming” Example

In order to provide a baseline for other tests, we selected a nominal set of parameters (G = 3, *r* = 2 mm, water, less than 10% fatigue) that best replicated the *in vivo* kinematics of the PL during frog swimming [22]. The PL muscle was maximally activated for 100 ms and the force produced was sufficient to drive the robotic limb through the water. The frog foot at the end of the robotic limb opened during limb protraction and closed during limb retraction. During the power stroke the muscle produced a peak force of approximately 9 N (Fig. 6 A1, black line). This resulted in a peak torque of 0.054 N-m at the robot motor to accelerate the limb. During muscle force production, the muscle shortened at a relatively constant rate (Fig. 6 A2). Force production ceased at 0.25 s at which point the muscle began to lengthen due to the force of gravity acting on the limb. Because of friction in the motor, the robotic limb did not completely return to its initial position (Fig. 6, row 2 and 3). In the nominal condition, the muscle produced 19 J of work per kilogram of muscle mass, as measured by the area enclosed by the work loop (Fig. 6 A3). The average power during the shortening phase was 77 W/Kg of muscle mass.

For each condition, we plotted muscle excursion versus muscle force to measure the work generated by the muscle. Because of the dynamics interactions between the muscle force and length, our technique differs importantly from classic work-loop techniques [13]. The term “work loop” typically refers to a particular set of classical experiments where the muscle length is oscillated in a predetermined trajectory that is independent of muscle force. Therefore, the work that is measured is only applicable to the conditions where forces from the environment and other muscles cause the muscle to move in the exact trajectory that is forced to move in. Here, changes in the work produced under similar muscle stimulation conditions are due to the coupled effects of muscle state on force production in response to identical muscle stimulation parameters, as well as the altered limb trajectory resulting from differences in muscle force production.

### B. Varying Moment Arm

To examine the role that biomechanical configuration can have on muscle work production, we compared two moment arm lengths (*r*): 2 mm (nominal condition) and 1 mm. Reducing the moment arm by one-half resulted in, one third of the work production compared to the nominal condition (Table III Biomechanics). Although the peak force produced by the muscle was greater using the shorter moment arm (1 mm), the torque applied to the robotic limb was approximately half of that applied during the nominal condition (2 mm). As a consequence, the robotic limb did not rotate as much through the water using the shorter moment arm. Due to combination of the shorter moment arm and the reduced rotation of the robotic limb, the muscle shortened at slower speed during the power stroke compared to the nominal condition. The decrease in work production using the shorter moment arm was primarily due to the 67% decrease in muscle length excursion.

### C. Varying Environment Viscosity

To illustrate how environmental viscosity can affect muscle work production, we allowed the robotic limb to rotate through air instead of water. A 1 mm moment arm was used because the nominal moment arm (2 mm) caused the robotic limb to rotate 360°. The work and average power generated by the muscle doubled when the limb rotated through air (Table III Environment) when compared to water. Although peak muscle force was similar in both conditions, total limb excursion was 3 times greater because of the decreased viscous resistance of air. In contrast to the other conditions, the muscle continued to shorten even after the muscle stopped producing a force. During protraction, the robotic device bounced off the mechanical stopper (indicated by the discontinuity in muscle length trajectory, Fig. 6 B2). The area enclosed by the work loop was 2.5 times greater when the viscosity of the environment was reduced.

### D. Effects of Fatigue

To demonstrate the possible effects of fatigue, we examined the force and work produced by fatigued muscles under the dynamic loading conditions; here our methods differ significantly from traditional work-loop approaches where the muscle length may be constrained to a nominal trajectory as muscle force decreases. We measure fatigue as the drop in muscle isometric force. With increasing muscle fatigue, the peak force produced and total length shortened during the power stroke declined (Fig. 6 C, Table III Fatigue). As peak force declined, less torque was applied to the robotic limb, causing both the total muscle excursion and the speed of shortening to decline (Fig. 6 C2). While decreased torque contributed to the decline in work production, the reduced speed of shortening further reduced work production, as evidenced by work loops that were triangular rather than rectangular in shape. For example at 48% fatigue (the isometric force generating capabilities of the muscle have dropped 48%), the work (force × length) produced declined by 71% and average power was reduced by 75%

## IV. DISCUSSION

We developed a closed-loop neuromechanical system that facilitates the study of interactions between an isolated muscle and environmental loads via a robotic device. Improving upon previous methods, which were limited to simple computational loads [17, 18], our system enables muscle kinematics and energetics to be studied under a variety of complex physical loads in a controlled manner that better mimics natural behavioral conditions. We were able to study a muscle interacting with the complex fluid dynamics of a frog foot in water, which could not have been accurately simulated computationally. The closed-loop neuromechanical system has the potential to improve our understanding of the dynamic interactions between a muscle and its environment that underlie natural movements, and could serve as a platform to test functional electrical stimulation (FES) methods for rehabilitation of movement.

Although we have just provided example data, our results suggest that the neuromechanical system can be used investigate muscles and their function under behaviorally-relevant dynamic conditions. We replicated muscle function in a simplified swimming task. Frog swimming is a complicated locomotor behavior that requires the coordination of multiple extensor and flexor muscles that interact with the environment via the frog foot [22, 23, 26-28]. The flexibility, multiple degrees of freedom, and asymmetrical movement of the frog foot create nonlinear hydrodynamics that are difficult to simulate in real-time. The muscle trajectories generated by our system produced features that are comparable to those found during natural frog swimming. In our nominal experimental condition, the change in muscle length was within 10% of that measured in *in vivo* during swimming in frogs; the duration of shortening was also within 20% of that measured *in vivo* [22]. During synchronous swimming when both hindlimbs move together, the plantaris longus muscle is not typically active during lengthening by antagonistic muscles [26]. Similarly, in all the conditions we tested, the muscle was not active and did not produce any force during the lengthening phase.

We demonstrated that our device can be useful to reveal the roles of muscle moment arm, environment viscosity, and fatigue during conditions that simulate the power stroke of swimming where the muscle starts from rest and then rapidly shortens. The test conditions demonstrate how our closed-loop coupling of a muscle to a physical device could be used to answer question that would note be possible using current *in vivo* and *in vitro* technologies:

### Moment Arm

Varying the biomechanical configuration of muscles may help us better understand the functional limits of muscles during movement. Changing the anatomy of muscles *in vivo* is prohibitive. Our method provides researchers with a tool to investigate the effects of musculoskeletal morphology on movement.

### Environmental Viscosity

Current closed-loop techniques, which do maintain system dynamics, are still unable to reproduce the complex environmental loads that occur *in vivo*. The closed-loop neuromechanical system, using a robotic and not a computational device, allows the systematic study of muscle under a variety of complex loads.

### Muscle Fatigue

Studying the mechanical properties of fatigued or injured muscle may help develop alternate strategies to improve function of impaired muscle. We illustrated how the closed-loop neuromechanical system allows the capabilities and contributions of fatigued muscle to movement to be accurately quantified. These results could not have been obtained using *in vivo* or traditional *in vitro* method because traditional work loop methods would require one to know in advance muscle fatigue or injury affects the movement.

In our neuromechanical system, the muscle trajectory is not prescribed, but is determined by the dynamic interactions between muscular and environmental forces. Unlike traditional isolated muscle protocols, which predetermine the length trajectory of muscle, our system dynamically couples an isolated muscle to physical load. Traditional muscle physiology methods were specifically designed to isolate and measure the individual fundamental properties of muscle. These properties are the basis of numerous mathematical models, which are used to predict the function of muscle during complex tasks. However, the emergent behavior of muscle that arises from the interactions of all its mechanical properties cannot be verified experimentally using classical methods. Therefore, the forward approach enabled by the closed-loop neuromechanical system allows the causal relationships between a muscle and its environment to vary, thus producing a range of different movement conditions. or fluid viscosity, can be assessed in a controlled fashion during causal, dynamic interactions. Recently developed closed-loop isolated muscle systems [17, 18] are also capable producing dynamic force-length relationships that are not prescribed. However, these systems use computational and not robotic devices, limiting their ability to reproduce the complex loads that occur in the natural environment.

Our approach could be extended to more complex experimental motor-control paradigms and robotic systems including those used to examine terrestrial locomotion [20] and balance [29], swimming [30], or flying [19]. Additionally, the robotic device does not need to be in the same physical location as the muscle apparatus. Through a remote connection it would be possible for the robotic device to be examined in different environments while leaving the muscle apparatus in the lab. The system can also be integrated with a diverse set of experimental test equipment that include different muscles and intact parts of nervous systems. Further, the architecture could be duplicated to include multiple muscles and robotic devices with multiple degrees of freedom.

Our closed-loop neuromechanical approach ultimately has the potential for application in clinical rehabilitation. Current FES research, which is largely concerned with minimizing muscle fatigue and increasing contraction force [31-33], may benefit from an improved understanding of fatigued muscle mechanics. Our system could be used to evaluate stimulation techniques [34, 35] on muscle — modified by physical injury, neural trauma, or fatigue — during interactions with complex environments. This technology may help advance our understanding of the neuromuscular system and help improve rehabilitation technologies.

## V. ACKNOWLEDGEMENTS

The authors would like to thank Edgar Brown for his technical advice and Brock Wester for his help with the illustrations. The project was funded in part by the Center for Behavioral Neuroscience (IBN-0349042) and the National Institute of Health (1 R01 EB006179-01A1).

